# From chambers to communities: gall architecture shapes parasitoid network stability in *Diplolepis* (Cynipidae, Diplolepididae) rose galls

**DOI:** 10.64898/2026.09.09.750361

**Authors:** Robert Veres, Zoltán László

**Affiliations:** Hungarian Department of Biology and Ecology, UBB Cluj, 400006, Romania; Centre 3B, Faculty of Biology and Geology, Babeș-Bolyai University, Cluj-Napoca, Romania

**Keywords:** connectance, *Diplolepis*, ecological network stability, gall architecture, parasitoid community, robustness

## Abstract

1. Gall-inducing insects create structurally distinct microhabitats that mediate their interactions with associated parasitoid communities, and variation in gall architecture is recognized as an important driver of the composition and organization of these communities.
2. In cynipid gall wasps of the genus *Diplolepis*, galls range from single-chambered to multichambered structures, and this variation has previously been linked to differences in parasitoid attack rates and community composition, yet its consequences for the topology and stability of the resulting ecological networks remain unexplored.
3. We compiled 17 empirical *Diplolepis*-parasitoid interaction networks spanning seven host taxa and used dynamical (non-normalized return time) and structural (robustness) measures of stability, together with network topology and diversity metrics, to test whether gall architecture shapes the organization of the resulting parasitoid networks. We hypothesized that networks associated with multichambered galls would be more stable, more connected and more unevenly structured than those associated with single-chambered galls.
4. Multichambered-gall networks showed significantly higher robustness at the species level and lower species-level non-normalized return times under intermediate and weak self-regulation, together with higher species-level connectance and clustering coefficients. In contrast, gall architecture reduced rather than increased Shannon diversity after accounting for network size, and had no detectable effect on the Gini index. Most effects weakened after aggregation to the genus level, highlighting the sensitivity of gall-associated network descriptors to taxonomic resolution.
5. These results indicate that gall architecture leaves a measurable signature on the organization and stability of associated parasitoid networks that extends beyond simple differences in community composition, adding a network-level perspective to the study of gall morphology as an ecological filter of natural-enemy communities.

## Introduction

Insect-induced plant galls are complex, insect-controlled modifications of host-plant tissue that provide developing larvae with both nutrition and a physical barrier against attack from natural enemies (Oliveira et al., 2016). Because parasitoids must nonetheless penetrate this barrier to reach the host, gall structure itself has long been considered a target of selection imposed by natural enemies, an idea most explicitly formalized as the Enemy Hypothesis (Stone & Schönrogge, 2003). Recent studies testing this hypothesis in several different gall systems show that both external and internal gall traits are linked to the type of enemies present in the community. Galls that place larvae farther from the surface tend to have more ectoparasitic, idiobiont enemies with long ovipositors (Baine et al., 2024). Complementary network-based approaches applied to galling insects and their parasitoids on shared host plants have likewise shown that morphological coupling between gall traits and parasitoid attack strategy, rather than host-plant identity alone, is the principal factor structuring these interactions (Luz et al., 2021; Prauchner & de Souza Mendonça, 2024). Gall morphology functions as an ecological filter that shapes not only which enemies attack a gall, but how those enemies are organized into a community.

Most evidence linking gall traits to natural-enemy communities has so far been framed in terms of species richness, composition or guild structure, rather than the organization of the interactions themselves. Ecological network approaches offer a complementary perspective, because metrics such as connectance, clustering and robustness summarize how interactions among community members are arranged and how that arrangement is expected to respond to perturbation, properties that are not captured by community composition alone. Explicitly network-based analyses of galling insects and their associated parasitoids and inquilines are comparatively scarce and recent (Fang et al., 2024), but the studies available indicate that host-plant identity, gall morphology and hybridization can all leave detectable signatures in the topology of the resulting galler-parasitoid networks (Luz et al., 2021; Prauchner & de Souza Mendonça, 2024). Whether structural variation in gall architecture translates into differences in network stability, as opposed to differences in topology or composition alone, has not been directly tested.

Ecological stability itself is a multidimensional property, encompassing distinct dynamical and structural components that need not respond in the same way to a given ecological driver (Donohue et al., 2016). Dynamical stability, typically summarized through the recovery time of a system following a small perturbation, depends strongly on the strength and distribution of self-regulation among interacting species, such that the same network topology can yield markedly different stability outcomes depending on how strongly species regulate their own dynamics (Barabás et al., 2017; Yang et al., 2023). Structural stability, in contrast, is more commonly assessed through robustness to sequential species loss and is closely tied to network connectance and the redundancy of trophic pathways (Allesina et al., 2009; Dunne et al., 2002; Gilbert, 2009). Because dynamical and structural stability capture different aspects of a network’s response to disturbance, and because taxonomic resolution itself can alter the network descriptors used to characterize antagonistic interactions (Llopis-Belenguer et al., 2023; Renaud et al., 2020; Rodrigues & Boscolo, 2020), a comprehensive test of whether gall architecture influences parasitoid network stability requires considering multiple stability measures across taxonomic resolutions rather than relying on any single metric.

Gall morphology can play an important role in shaping the communities associated with gall-inducing insects. Because parasitoids must interact with their hosts through the gall tissues, structural characteristics of the gall can influence both host accessibility and parasitoid community composition. In cynipid gall systems, traits of the gall and the spatiotemporal niche of the host have been shown to explain substantial variation in parasitoid community structure, with host aggregation and multilocularity also contributing to differences among parasitoid assemblages (Bailey et al., 2009). Similar patterns have been demonstrated specifically in the case of *Diplolepis*, where gall size, chamber number and chamber-wall characteristics were associated with parasitoid incidence and attack rates, and differences in gall morphology were suggested to contribute to differences in the associated parasitoid communities (László & Tóthmérész, 2013). Multilocular *D. rosae* galls can also support complex internal communities, in which changes caused by inquilines can significantly alter the number and diversity of emerging insects (László & Tóthmérész, 2006). More recent comparisons across several gall-inducing insect communities, including *Diplolepis*, further demonstrate that variation in gall morphology is associated with the functional composition of natural-enemy communities (Baine et al., 2024).

Differences between single- and multichambered galls may therefore extend beyond parasitoid species composition and be reflected in the organization of the resulting ecological networks. The presence of multiple larval chambers within the same gall provides greater opportunities for different parasitoid species to occur within the same local community, potentially increasing the number of realized associations among its members. Consequently, networks associated with multichambered galls can be expected to be more highly connected than those associated with single-chambered galls. Network topology, in turn, can strongly influence responses to disturbance. Higher connectance has been associated with increased robustness against secondary extinctions in empirical food webs (Dunne et al., 2002), while heterogeneity in the distribution of interactions among species can also influence dynamical stability, although its effects depend on the type and organization of ecological interactions (Yan et al., 2017). We therefore expected the differences in the spatial and biological structure of single- and multichambered galls to be reflected in both the stability and topology of their associated parasitoid networks.

We hypothesized that parasitoid networks associated with multichambered *Diplolepis* galls are ecologically more stable than those associated with single-chambered galls. Specifically, we predicted that networks from multichambered galls would exhibit smaller return time (RT) values, indicating greater dynamical stability following perturbation, and higher robustness (R), indicating greater structural stability against species loss. We further hypothesized that multichambered-gall networks would have higher connectance (C), which indicates a greater proportion of realized interactions among network members, and higher Gini index (G) values, indicating a more uneven distribution of interactions among species.

## Materials and Methods

A comprehensive search of the literature resulted in 12 empirical parasitoid communities containing sufficient information to properly reconstruct interaction networks. These were completed with 5 unpublished communities based on our collecting and rearing. The obtained 17 communities consisted of seven *Diplolepis* taxa: *D. abei* (n = 1), *D. eglanteriae* (n = 2), *D. fructuum* (n = 3), *D. nervosa* (n = 2), *D. spinosissimae* (n = 2), *D. rosae* (n = 5), and *D. mayri* (n = 2) (Supp. Table 1).

Community interaction matrices were generated separately at the species and genus levels using the abundance data available for each community. In the species-level networks, taxa identified only to genus in the original data were retained as separate genus-level nodes rather than being excluded or assigned to species without supporting information, thereby preserving their observed abundances and trophic interactions. These records represented 4 of the 31 associate taxa and approximately 1.7% of all recorded associate individuals (Supp. Table 1). For simplicity, these two representations are hereafter referred to as the species-level and genus-level networks, respectively.

Gall-inducer abundances were not available for the three *D. fructuum* communities and were estimated from the total abundance of associated taxa using a gall-inducer-to-associate ratio of 0.53, derived from the seven *D. rosae* and *D. mayri* communities for which gall-inducer abundance was known (Supp. Table 1). To assess the sensitivity of the results to the estimated gall-inducer abundances, all statistical models were additionally rerun after excluding the three *D. fructuum* networks for which gall-inducer abundance had been estimated. Observed abundances were subsequently allocated among interacting taxa according to a predefined set of biologically permitted trophic relationships (Supp. Table 1). For taxa known to act both as primary parasitoids and hyperparasitoids, abundance was divided equally between interactions with the focal gall inducer and their corresponding hyperparasitic interactions. In contrast, the abundance of taxa acting exclusively as primary parasitoids or exclusively as hyperparasitoids was assigned entirely to the corresponding interaction (Supp. Table 1). The complete set of 17 community matrices used in subsequent analyses is provided in Supp. Table 2.

Network stability was assessed from both dynamical and structural perspectives using non-normalized return times (RT) and robustness (R), respectively, following the general methodology of Veres and László (2026). All stability analyses were conducted separately at both the species and genus levels. Non-normalized return times were calculated from the leading eigenvalue of the parametrized community matrices and represent the characteristic recovery times of locally stable systems following a small perturbation. For matrices in which the leading eigenvalue had a negative real part, RT was calculated as RT = −1/Re(λ_max_), where Re(λ_max_) < 0. Lower RT values therefore indicate faster recovery and greater dynamical stability; because interaction strengths are row-standardized proportions rather than measured rates, RT is a relative, dimensionless index rather than a return time in real (e.g., generational) time units. In contrast to Veres and László (2026), where interaction strengths were derived from a common set of fixed starting coefficients, the initial interaction strengths used in the present study were calculated individually for each interaction prior to randomization.

Interaction strengths were calculated from the raw abundance matrices. First, main diagonal values were set to zero, after which each interaction count was divided by the total count of its corresponding row. The resulting values represented the proportion of a taxon’s recorded interactions attributable to each interaction partner. Row-standardized values above the main diagonal were retained as positive effects, whereas corresponding values below the diagonal were assigned a negative sign to represent effects in the opposite direction; absent interactions remained zero. Self-regulation was subsequently introduced along the main diagonal using constant values of −1, −0.5, or −0.15, chosen to represent progressively weaker negative self-effects. This range was used to assess the sensitivity of network dynamics to self-regulation strength, motivated by theoretical evidence that the magnitude of negative self-effects can strongly influence local stability (Barabás et al., 2017). Alternatively, a taxon-specific self-regulation value was calculated from each taxon’s total abundance in the original matrix. Under this final, abundance-dependent parameterization, self-regulation became increasingly negative with increasing taxon abundance and approached −1 at high abundances.

Monte Carlo simulations were performed separately for each of the 17 communities, four self-regulation parameterizations, and two taxonomic resolutions, resulting in 136 simulation sets. Each simulation set consisted of 1,000,000 repetitions during which every non-zero starting interaction strength was independently multiplied by a random value drawn from a U(0,1) distribution before calculating the leading eigenvalue of the resulting community matrix (Pimm & Lawton, 1977; Veres & László, 2026). Only simulations producing a leading eigenvalue with a negative real part, representing locally stable systems, were retained for the RT analysis. Non-normalized RT values were used throughout the analysis because the seasonal nature of all examined gall communities places them on comparable biological time scales, rendering the time-scale standardization applied in Veres and László (2026) unnecessary. For each simulation set, the retained RT distribution was condensed into a single representative non-normalized RT value based on its modal region, following the frequency-distribution procedure described in Veres and László (2026). For the abundance-dependent parameterization, two networks with extreme RT values (Dspinosissimae_13 and Dfructuum_16) were excluded from the statistical model because their inclusion resulted in a singular model fit.

Structural stability was quantified using network robustness based on simulated secondary-extinction sequences. Taxa were progressively removed from the network, and the resulting loss of dependent taxa was recorded to generate an attack-tolerance curve. Robustness was calculated as the normalized area under this curve, with values approaching 0 indicating structurally fragile networks and values approaching 1 indicating networks that tolerate a greater proportion of species loss before collapsing. Robustness simulations were repeated 1,000 times for each network across the two taxonomic resolutions, and the mean robustness across repetitions was used in subsequent analyses. The calculation and interpretation of robustness followed Veres and László (2026).

In addition to non-normalized return time and robustness, several metrics describing network topology and community diversity were calculated for each network. Connectance was used to quantify the proportion of realized interactions within a community and was calculated as *C* = 2*L*/[*S*(*S* - 1)], where *L* represents the number of realized interactions and *S* the number of taxa in the network. The clustering coefficient was used to describe the tendency of interacting taxa to form locally interconnected groups and was calculated as the mean local clustering coefficient of the network. For each taxon *i*, the local clustering coefficient was defined as the proportion of realized links among its *k_i_* neighbors relative to the *k_i_*(*k_i_* - 1)/2 possible links among those neighbors; taxa with fewer than two neighbors were assigned a value of zero. Shannon diversity was calculated from taxon abundances using the ‘vegan’ package (Oksanen et al., 2001). All metrics were calculated separately for the species- and genus-level networks.

Inequality in the distribution of interactions among taxa was quantified with the Gini index, calculated using the ‘igraph’ (Csárdi et al., 2006) and ‘ineq’ (Zeileis, 2000) packages. For each weighted network, taxon strengths were calculated using igraph::strength(), and the Gini index of the resulting strength distribution was obtained using ineq::ineq(…, type = “Gini”), with higher values indicating a more uneven distribution of interaction strength among network members. Network modularity was additionally quantified using the weighted Leiden community detection implemented in the ‘igraph’ package. Communities were identified using igraph::cluster_leiden() with interaction strengths supplied as edge weights and a resolution parameter of 1. The resulting community membership was then used to calculate weighted modularity with igraph::modularity().

Both Gini index and modularity calculations were based on undirected weighted graphs constructed from the raw interaction-count matrices, with observed interaction counts used as edge weights. At the genus level, two networks with extreme modularity values (Dfructuum_2 and Dnervosa_15) were excluded from the statistical model because their inclusion resulted in a singular model fit.

Statistical analyses were performed using linear mixed-effects models implemented in the ‘nlme’ (Pinheiro & Bates, 1999) package. For each network metric, the general model structure was *metric ∼ network size (β_1_) + gall architecture (β_2_)*. Network size and gall architecture were included as fixed effects, and *gall architecture* distinguished between single- and multichambered galls, with the single-chambered level serving as baseline, and network identity was included as a random effect in all linear mixed-effects models. For non-normalized return times, models were fitted separately to each combination of the four self-regulation parameterizations and two taxonomic resolutions, resulting in eight models. For all other network metrics, separate models were fitted at the species and genus levels, resulting in two models per metric.

All statistical metrics calculated for the 17 communities and used in subsequent analyses are available in Supp. Table 3.

All statistical analyses were performed in the R v4.6.0 statistical environment (R Core Team, 2026). General data handling and manipulation were performed using the ‘tidyverse’ (Wickham, 2016) collection of packages. Statistical figures were produced using ‘ggplot2’ (Wickham et al., 2007), while the interaction structure of the analyzed communities was additionally visualized as Sankey diagrams using ‘plotly’ (Sievert et al., 2015). Further statistical model metrics were calculated using ‘broom.mixed’ (Bolker & Robinson, 2018) and ‘emmeans’ (Lenth & Piaskowski, 2017) packages.

## Results

The dataset comprised 17 analyzed *Diplolepis* parasitoid networks, including 7 communities associated with single-chambered galls and 10 communities associated with multichambered galls. At the species level, network size ranged between 6 - 16 taxa (mean = 9.71, median = 10), with 6 - 19 realized interactions per network (mean = 11.06, median = 11). Aggregation to the genus level reduced network size to 6 - 12 genera (mean = 8.41, median = 8), with 6 - 12 realized interactions per network. Multichambered-gall communities were larger on average than single-chambered communities at both taxonomic resolutions. The difference in network size between gall architectures was marginally significant at the species level (mean = 10.80 vs. 8.14; β = −2.66, SE = 1.37, t = −1.94, p = 0.072) and significant at the genus level (mean = 9.20 vs. 7.29; β = −1.91, SE = 0.79, t = −2.41, p = 0.029). The taxonomic composition, interaction structure and abundances of the analyzed communities are summarized in Fig. 1.

**Figure 1.**
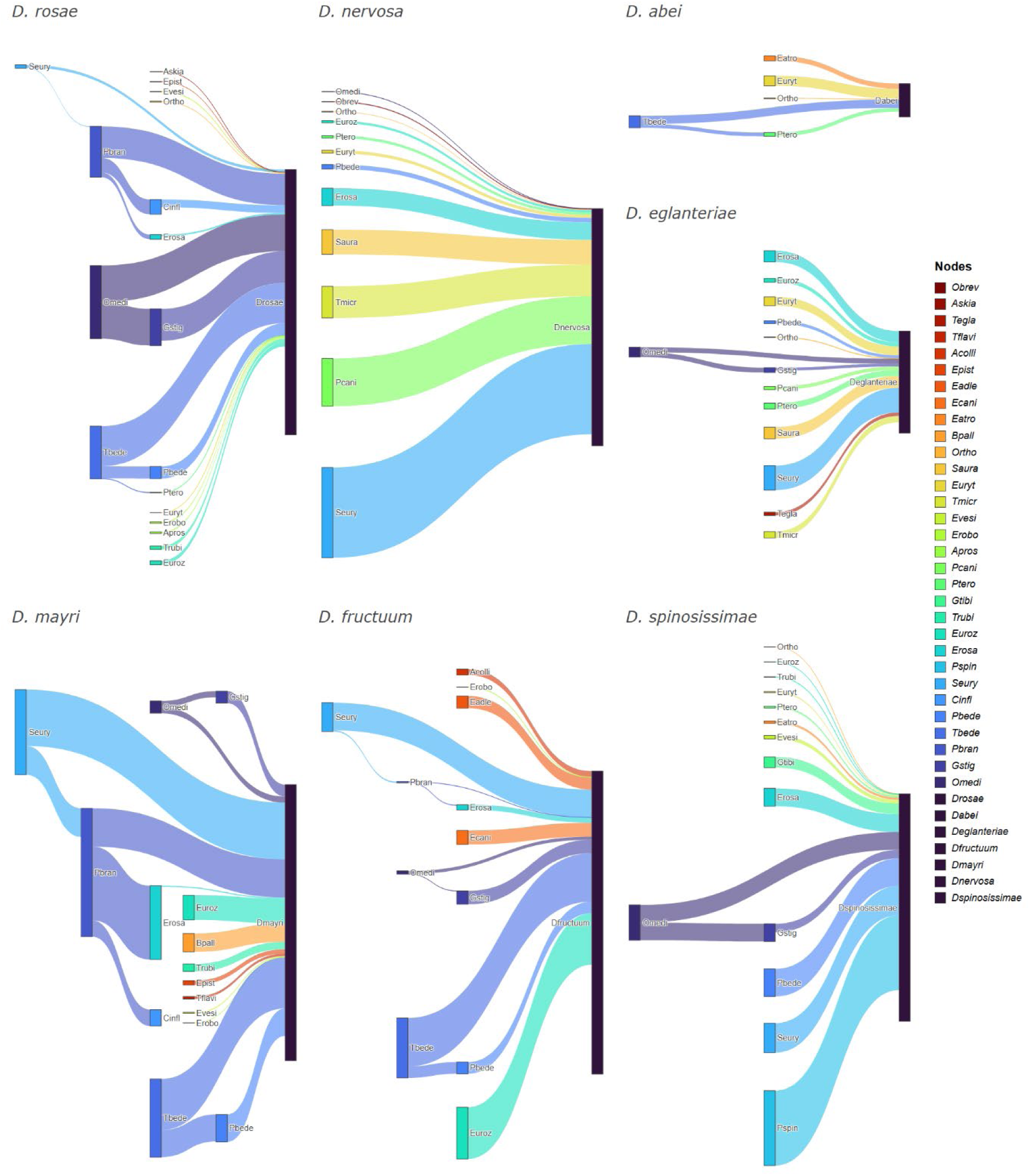
Sankey diagrams showing the reconstructed parasitoid interaction networks associated with the seven *Diplolepis* taxa. For clarity, networks were merged by gall inducer, such that all interactions recorded across replicate networks of the same *Diplolepis* taxon were combined into a single diagram. The figure shows one summarized network for each gall inducer rather than the full set of 17 individual networks. The flow widths are proportional to the combined interaction abundances. The unlinked lower portion of each gall-inducer node represents individuals that emerged successfully without being parasitized. Taxa identified only to genus in the original data are shown at the genus level, namely ‘Ortho’, ‘Ptero’, ‘Euryt’, and ‘Apros’. *D. abei*, *D. eglanteriae*, *D. nervosa*, and *D. spinosissimae* were recorded as single-chambered taxa, whereas *D. fructuum*, *D. mayri*, and *D. rosae* were recorded as multichambered taxa.

**Figure 2.**
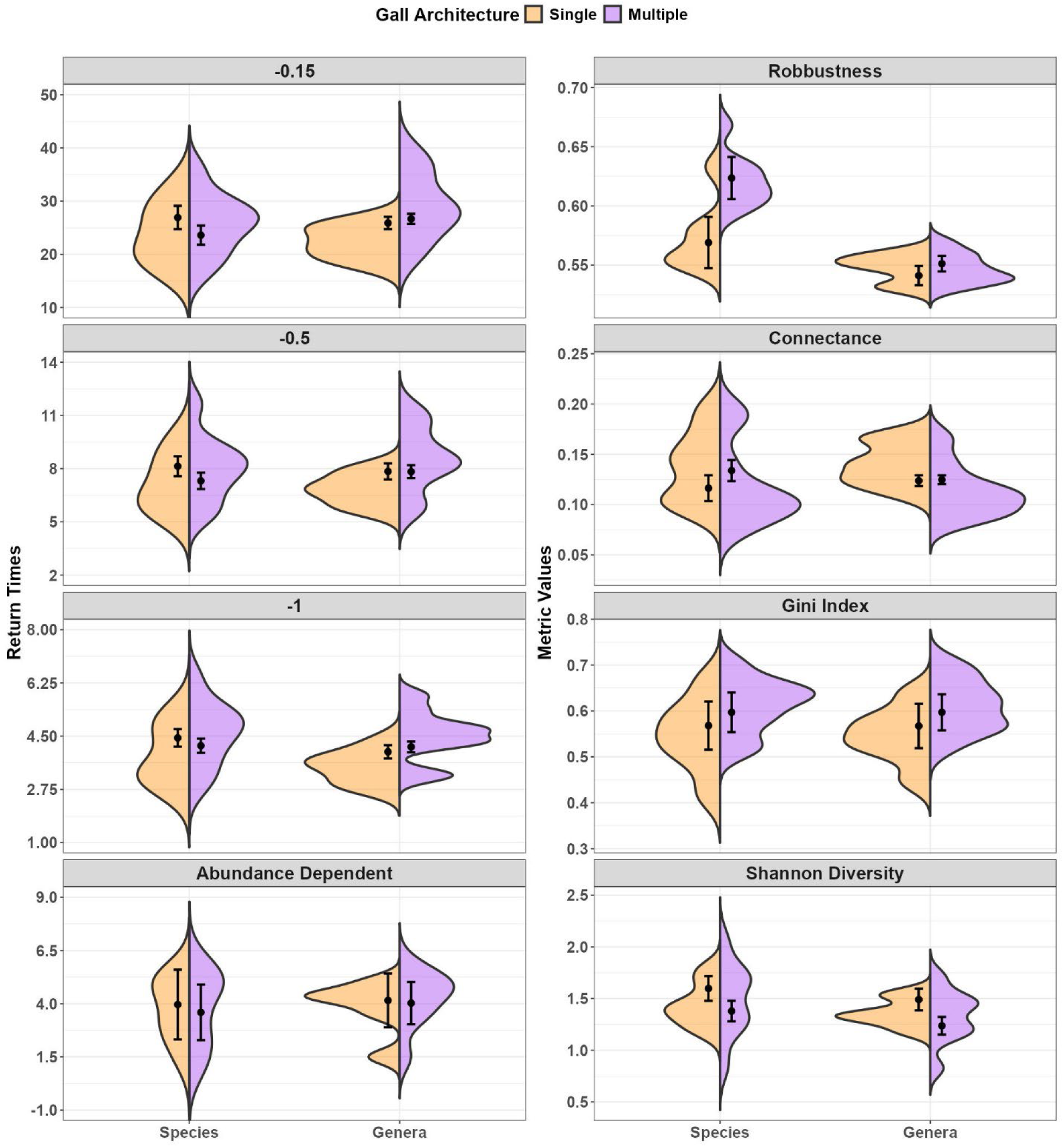
Comparison of stability, network-topological, and diversity metrics between parasitoid communities associated with single- and multichambered *Diplolepis* galls at species and genus taxonomic resolutions. Each panel shows differences between gall architectures (single-chambered and multichambered) and between species- and genus-level representations of the same networks. The four panels on the left show non-normalized return times (RT) obtained under the three constant self-regulation parameterizations (−0.15, −0.5, and −1) and the abundance-dependent self-regulation parameterization. The four panels on the right show robustness, connectance, Gini index, and Shannon diversity. Violin plots show the comparison and distribution of metric values among the analyzed networks. Black points and error bars show the estimated marginal means and 95% confidence intervals obtained from the linear mixed-effects models after accounting for network size. The distributions shown here correspond to the network-level values used in the linear mixed-effects models evaluating the effects of network size and gall architecture reported throughout the Results and in Table 1.

**Figure 3.**
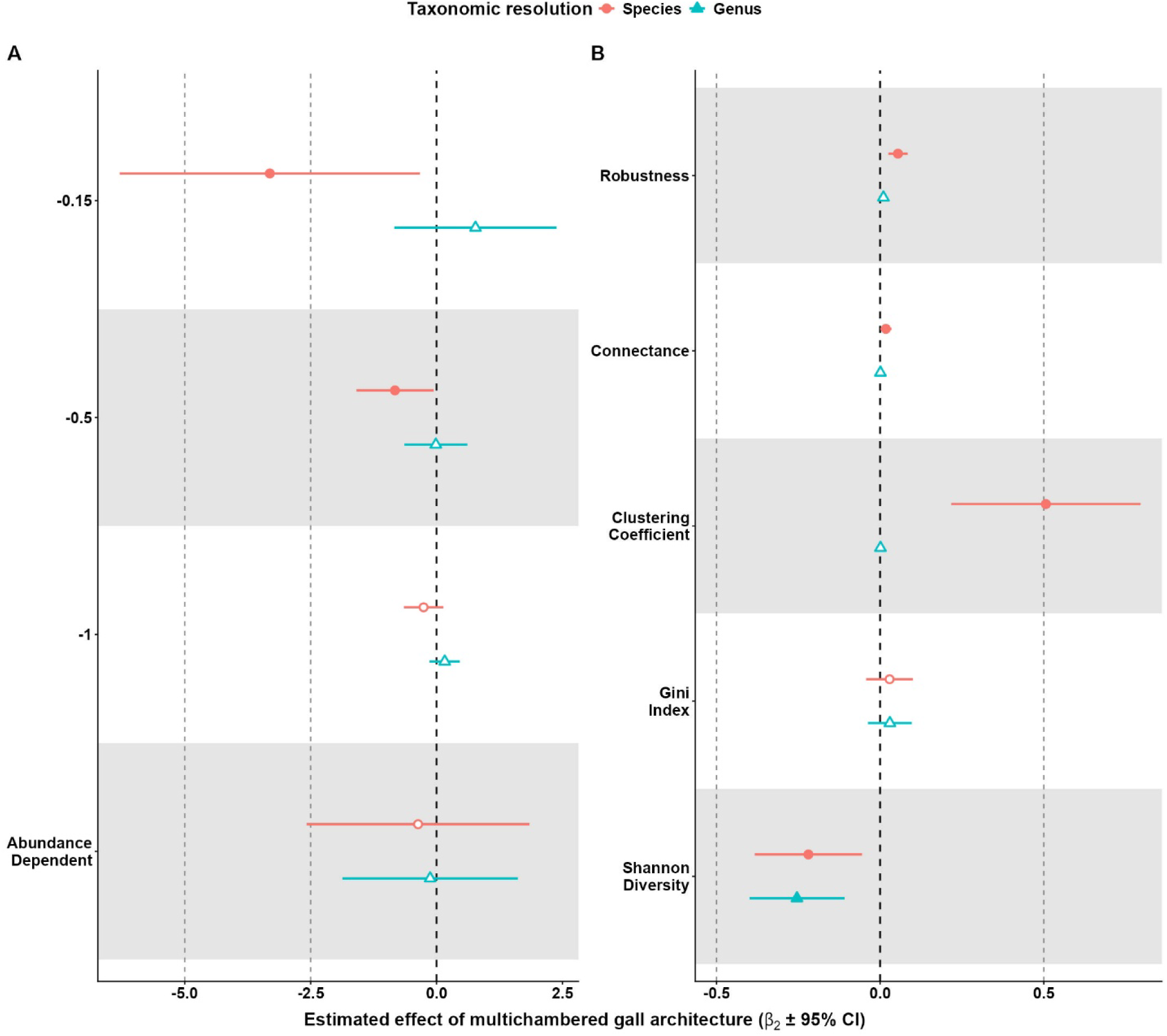
Forest plot of the estimated effects of multichambered gall architecture on parasitoid network metrics at species- and genus-level taxonomic resolutions. Effect sizes are shown as the model coefficients for multichambered relative to single-chambered galls (β_2_) with 95% confidence intervals from the linear mixed-effects models. **(A)** Effects of gall architecture on non-normalized return time (RT) under the four self-regulation parameterizations: constant diagonal values of - 0.15, −0.5, and −1, and the abundance-dependent parameterization. **(B)** Effects of gall architecture on robustness, connectance, clustering coefficient, Gini index, and Shannon diversity. Filled symbols indicate significant effects (p < 0.05), whereas open symbols indicate non-significant effects. The vertical dashed line at zero denotes no difference between single- and multichambered galls; values to the right indicate higher metric values in multichambered networks, whereas values to the left indicate lower values. For RT, negative coefficients indicate lower return times in multichambered networks and therefore greater dynamical stability relative to single-chambered networks. All estimates are adjusted for network size.

The majority of Monte Carlo simulations produced locally stable community matrices from which RT values could be calculated. At the species level, the mean numbers of locally stable simulations out of 1,000,000 repetitions were 999,152 for a constant self-regulation of −1, 996,543 for −0.5, 966,722 for −0.15, and 936,922 under the abundance-dependent parameterization. The latter also showed the greatest variation among networks, with the number of stable simulations ranging from 498,521 to 1,000,000. At the genus level, all 1,000,000 simulations were locally stable for each network under the three constant self-regulation parameterizations, whereas the abundance-dependent parameterization produced a mean of 926,288 stable simulations, ranging from 249,741 to 1,000,000. Compared with the −1 self-regulation parameterization, the abundance-dependent parameterization produced significantly fewer locally stable simulations at both the species (β = - 62,230.6, SE = 29,374.5, t = −2.12, p = 0.039) and genus levels (β = −73,711.8, SE = 36,465.4, t = −2.02, p = 0.049). Gall architecture did not significantly affect the number of locally stable simulations at either the species (β = −8,811.2, p = 0.682) or genus level (β = −797.9, p = 0.977).

Non-normalized return times were strongly associated with network size under most self-regulation parameterizations, whereas the effect of gall architecture was dependent on both taxonomic resolution and self-regulation strength. At the species level, network size had a significant positive effect on RT when constant diagonal values of −1, −0.5, and −0.15 were used (β_1_ = 0.35 - 1.85, all p < 0.001), indicating increasing non-normalized return times with increasing network size. No significant relationship was found with network size under the abundance-dependent self-regulation parameterization (β_1_ = 0.11, p = 0.53, n = 15). Multichambered-gall networks exhibited significantly lower RT values than single-chambered networks when self-regulation was set to −0.5 (β_2_ = −0.83, SE = 0.36, t = −2.31, p = 0.037) and −0.15 (β_2_ = −3.31, SE = 1.39, t = −2.38, p = 0.032), supporting greater dynamical stability of multichambered networks under these parameterizations.

Gall architecture had no significant effect when the diagonal was set to −1 (β_2_ = −0.26, p = 0.181) or when abundance-dependent self-regulation was used (β_2_ = −0.366, p = 0.725, n = 15).

At the genus level, network size likewise had a significant positive relationship with RT under all three constant self-regulation parameterizations (β_1_ = 0.43 - 3.12, all p < 0.001). Under the abundance-dependent parameterization, the relationship was not significant (β_1_ = 0.26, p = 0.28, n = 15). Gall architecture did not significantly affect RT under any of the four genus-level parameterizations (all p ≥ 0.266). Full model estimates for the effects of network size and gall architecture across all four self-regulation parameterizations and both taxonomic resolutions are presented in Table 1.

**Table 1.**
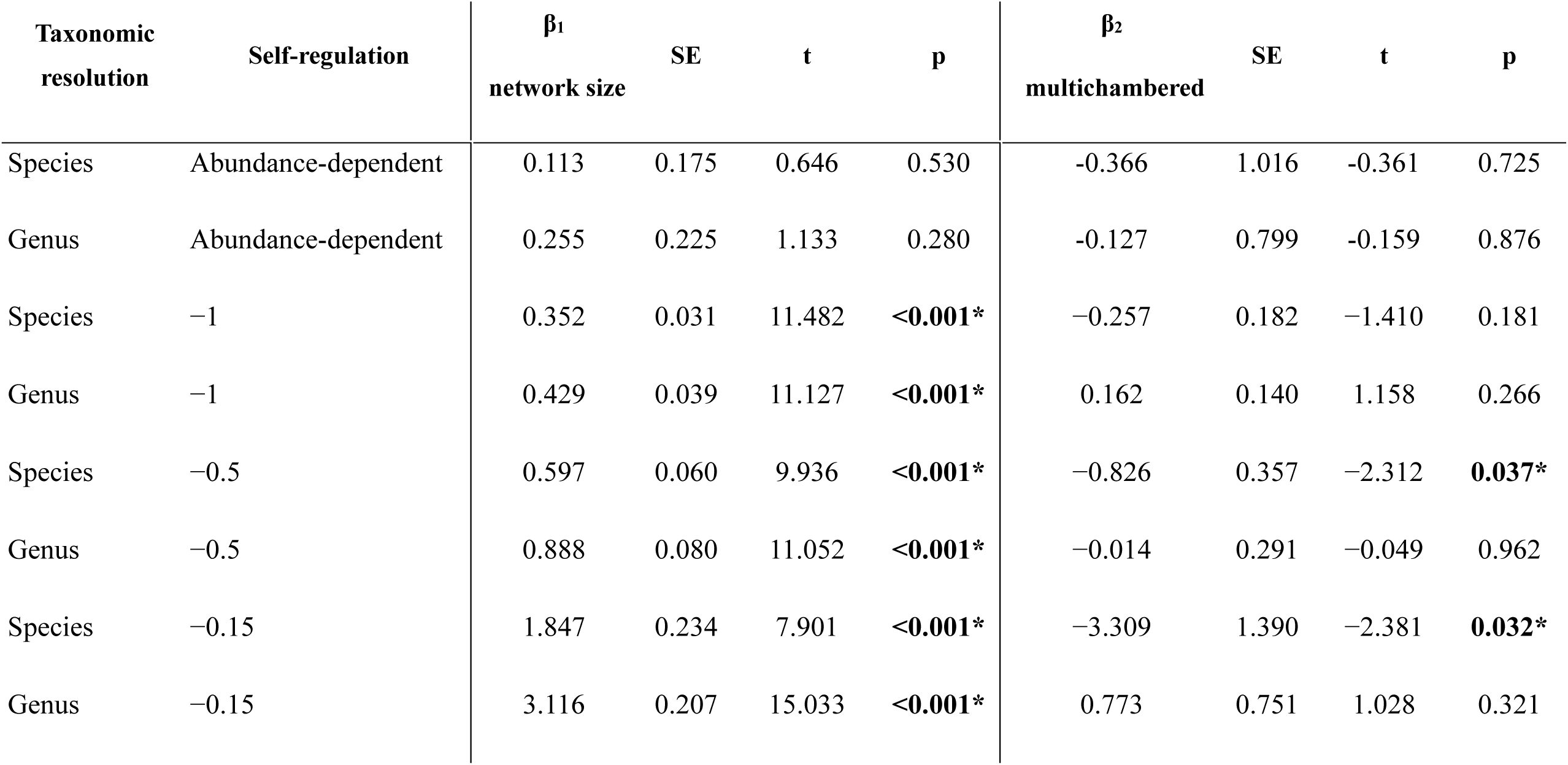
Linear mixed-effects model results for non-normalized return times across four self-regulation parameterizations and two taxonomic resolutions. Network identity was included as a random effect. The coefficient for gall architecture represents the effect of multichambered relative to single-chambered galls; consequently, negative coefficients indicate lower non-normalized return times in multichambered networks. Abundance-dependent models were based on 15 networks at each taxonomic resolution; constant self-regulation models were based on all 17 networks.

Multichambered-gall communities showed higher estimated robustness at both taxonomic resolutions, with the effect significant at the species level and marginally significant at the genus level. At the species level, gall architecture had a significant positive effect on robustness (β = 0.055, SE = 0.014, t = 3.97, p = 0.001), whereas network size had no significant effect (β = −0.002, SE = 0.002, t = −1.07, p = 0.302). At the genus level, the effect of gall architecture remained positive but was marginally significant (β = 0.010, SE = 0.005, t = 1.94, p = 0.073). Network size had a significant negative effect on genus-level robustness (β = −0.005, SE = 0.001, t = −3.82, p = 0.002), indicating decreasing robustness with increasing network size after accounting for gall architecture.

Overall, the stability analyses provided partial support for the prediction that multichambered-gall parasitoid networks are more stable, with the clearest and most consistent support arising from structural robustness and from species-level RT under the −0.5 and −0.15 self-regulation parameterizations.

Connectance decreased significantly with increasing network size at both taxonomic resolutions (species: β_1_ = −0.013, SE = 0.001, t = −9.50, p < 0.001; genus: β_1_ = −0.014, SE = 0.001, t = −15.14, p < 0.001). At the species level, multichambered-gall networks had significantly higher connectance than single-chambered networks (β_2_ = 0.017, SE = 0.008, t = 2.16, p = 0.049). At the genus level, however, gall architecture had no significant effect on connectance (β_2_ = 0.001, SE = 0.003, t = 0.28, p = 0.781).

The clustering coefficient showed a similar dependence on taxonomic resolution. At the species level, multichambered-gall networks had significantly higher clustering coefficients than single-chambered networks (β_2_ = 0.506, SE = 0.135, t = 3.75, p = 0.002), whereas network size had no significant effect (β_1_ = −0.011, SE = 0.023, t = −0.49, p = 0.630). At the genus level, gall architecture had no significant effect on clustering coefficient (β_2_ = 0.001, SE = 0.003, t = 0.34, p = 0.742), while clustering decreased significantly with increasing network size (β_1_ = −0.009, SE = 0.001, t = −9.60, p < 0.001).

The Gini index increased with network size at both taxonomic resolutions, although the strength of this relationship differed between them. At the species level, network size had a significant positive effect on Gini index values (β_1_ = 0.013, SE = 0.006, t = 2.29, p = 0.038), whereas at the genus level the positive relationship was marginally significant (β_1_ = 0.018, SE = 0.009, t = 2.07, p = 0.057). Multichambered-gall networks also had slightly higher estimated Gini index values than single-chambered networks at both resolutions, but the effect of gall architecture was not significant at either the species (β_2_ = 0.029, SE = 0.033, t = 0.87, p = 0.399) or genus level (β_2_ = 0.030, SE = 0.031, t = 0.96, p = 0.355).

Shannon diversity increased significantly with network size at both the species (β_1_ = 0.097, SE = 0.013, t = 7.51, p < 0.001) and genus levels (β_1_ = 0.112, SE = 0.019, t = 6.00, p < 0.001). In contrast, gall architecture had a significant negative effect on Shannon diversity at both taxonomic resolutions, with multichambered-gall communities having lower diversity after accounting for network size (species: β_2_ = −0.219, SE = 0.076, t = −2.87, p = 0.012; genus: β_2_ = −0.254, SE = 0.068, t = −3.75, p = 0.002).

Modularity was not significantly related to either network size or gall architecture at either taxonomic resolution. At the species level, neither network size (β_1_ = 0.0008, SE = 0.0016, t = 0.50, p = 0.627) nor gall architecture (β_2_ = 0.0080, SE = 0.0097, t = 0.82, p = 0.424) had a significant effect on modularity. At the genus level, neither network size (β_1_ = 0.00001, SE = 0.00006, t = 0.18, p = 0.863, n = 15) nor gall architecture (β_2_ = 0.00027, SE = 0.00022, t = 1.21, p = 0.251, n = 15) significantly affected modularity.

For the aforementioned sensitivity analysis, excluding the three *D. fructuum* networks for which gall-inducer abundances were estimated produced little qualitative change in the model results, with 20 of the 22 relevant models retaining the same significance pattern as in the full dataset.

The only changes were that the species-level effect of gall architecture on connectance and the species-level effect of network size on the Gini index both became non-significant after the exclusion.

## Discussion

Our results provide support for the hypothesis that parasitoid networks associated with multichambered *Diplolepis* galls are more stable than those associated with single-chambered galls. The clearest support was provided by structural stability, with multichambered networks showed significantly higher robustness at the species level, while the same positive relationship was marginally significant at the genus level. The difference between the responses of structural and dynamical stability is consistent with the multidimensional nature of ecological stability, whereby different stability properties can respond differently to the same characteristics of an ecological community (Donohue et al., 2016). Dynamical stability showed a more conditional response. Multichambered networks had significantly lower return times at the species level when self-regulation was set to −0.5 and −0.15, whereas no significant difference was observed under the −1 or abundance-dependent parameterizations, and no significant effect of gall architecture was detected at the genus level. Such dependence of dynamical stability on self-regulation is consistent with theoretical work demonstrating that the magnitude and distribution of negative self-effects can substantially influence the local stability of ecological networks (Barabás et al., 2017; Yang et al., 2023). Gall architecture was also associated with differences in network topology, with multichambered networks showing higher connectance and clustering coefficients at the species level. Morphological characteristics of galls have previously been shown to structure parasitoid communities and, more specifically, galler-parasitoid interaction networks (Bailey et al., 2009; Baine et al., 2024; Luz et al., 2021; Prauchner & de Souza Mendonça, 2024). Together, these results suggest that gall architecture can influence not only the composition of the associated parasitoid community, but also the organization and stability of the resulting ecological network (Bailey et al., 2009; Baine et al., 2024; Luz et al., 2021; Stone & Schönrogge, 2003).

The association between gall architecture and parasitoid network structure is consistent with previous evidence that gall morphology acts as an important ecological filter of natural-enemy communities (Bailey et al., 2009; Baine et al., 2024; Joseph et al., 2011; Stone & Schönrogge, 2003). Structural gall traits have been shown to influence parasitoid access to concealed hosts and have been associated with substantial variation in parasitoid-community composition (Bailey et al., 2009; Joseph et al., 2011; Stone & Schönrogge, 2003). Within *Diplolepis*, László and Tóthmérész (2013) demonstrated that chamber number, gall size and chamber-wall characteristics are associated with parasitoid incidence and attack rates, and proposed that morphological differences between galls may contribute to differences among their parasitoid communities. More recently, gall morphology has also been shown to be associated with the functional composition of natural-enemy communities across several gall systems, including *Diplolepis* (Baine et al., 2024). Our results extend these patterns from parasitoid occurrence and community composition to the level of network organization, consistent with evidence that morphological coupling between gallers and parasitoids can contribute directly to the structure of their interaction networks (Luz et al., 2021; Prauchner & de Souza Mendonça, 2024). The greater species-level connectance and clustering observed in multichambered galls indicate that differences in gall structure can be reflected in the way parasitoid-community members are linked to one another. The presence of multiple larval chambers may provide greater opportunities for different parasitoid and hyperparasitoid taxa to occur within the same gall system, thereby allowing a larger and more locally interconnected interaction network to develop (Bailey et al., 2009; Joseph et al., 2011; László & Tóthmérész, 2013).

The higher robustness of multichambered networks provides the strongest evidence that these differences in network organization have consequences for ecological stability. Robustness describes the tolerance of a network to species loss and the resulting secondary-extinction cascades, and has been widely used to characterize the structural response of food webs to extinction sequences (Allesina et al., 2009; Dunne et al., 2002, 2004). Consequently, the higher values observed in multichambered networks indicate that a greater proportion of species can be removed before substantial network collapse occurs. Food-web robustness has repeatedly been associated with network connectance and the organization of trophic links (Allesina et al., 2009; Dunne et al., 2002, 2004; Gilbert, 2009). Dunne et al. (2002) specifically showed that more highly connected food webs generally tolerate species loss better before extensive secondary extinctions occur. The simultaneous increase in connectance and robustness observed in the present study is consistent with this established relationship. Greater connectivity may provide greater redundancy in trophic pathways and reduce dependence on individual interaction partners, thereby increasing tolerance to the loss of particular species (Allesina et al., 2009; Dunne et al., 2002). However, because the present analyses did not explicitly test whether connectance mediates the effect of gall architecture on robustness, this relationship should be interpreted as an association rather than evidence for a direct causal mechanism.

The return-time analyses indicate that the effect of gall architecture on dynamical stability depends on the assumed strength of self-regulation. Under the three constant self-regulation parameterizations, non-normalized return times increased significantly with network size at both taxonomic resolutions. However, a significant difference between gall architectures was detected only at the species level when self-regulation was set to −0.5 and −0.15, in both cases with multichambered networks having lower return times and therefore faster recovery following perturbation. No gall-architecture effect was detected when self-regulation was set to −1, while the abundance-dependent parameterization produced neither a significant architecture effect nor the strong positive relationship between network size and RT observed under the constant parameterizations. Self-regulation is known to play a major role in determining the local stability of ecological community matrices, with negative self-effects capable of strongly influencing whether perturbations decay through time (Barabás et al., 2017; Yang et al., 2023). Variation among species in self-regulation strength can itself alter the dynamical response of ecological communities to perturbation (Barabás et al., 2017; Yang et al., 2023). The absence of a detectable gall-architecture effect under the strongest constant self-regulation may therefore indicate that strong negative self-effects can dominate the contribution of differences in interspecific network structure (Barabás et al., 2017). Conversely, under intermediate and weaker self-regulation, structural differences between single- and multichambered communities may contribute more strongly to their recovery dynamics. This interpretation is also consistent with the lower number of locally stable realizations produced by the abundance-dependent parameterization, indicating that assumptions concerning intraspecific regulation substantially influence the dynamical behavior of these small parasitoid networks.

Network size itself emerged as an important determinant of several network properties. Under all three constant self-regulation parameterizations, larger networks had longer return times, while connectance declined strongly with increasing network size at both species and genus levels. Genus-level robustness also decreased with increasing network size. These patterns are broadly consistent with classical expectations for relatively small ecological systems, in which increasing system size and complexity can reduce local dynamical stability (May, 1972, 2001; Pimm & Lawton, 1977, 1978; Veres & László, 2026). May’s theoretical work demonstrated that increasing the number of interacting components can reduce the probability of stability in randomly assembled ecological systems, while Pimm and Lawton similarly linked trophic complexity to slower recovery dynamics in small ecological communities (May, 1972, 2001; Pimm, 1982; Pimm & Lawton, 1977, 1978). Subsequent theoretical work has shown, however, that the relationship between complexity and stability also depends strongly on non-random properties of ecological network structure, including degree distributions, intervality and correlations among interactions (Allesina et al., 2015). The *Diplolepis* networks examined here fall entirely within a small network-size range, making these classical small-system predictions especially relevant. Importantly, however, gall architecture retained significant effects on several metrics even after network size was included in the models. Thus, the differences observed between single- and multichambered communities cannot be attributed solely to the generally larger size of multichambered networks.

The differences between species- and genus-level analyses further demonstrate that taxonomic resolution can substantially affect the ecological patterns detected in these networks, as has been shown for several types of ecological interaction networks (Llopis-Belenguer et al., 2023; Renaud et al., 2020; Rodrigues & Boscolo, 2020). Gall architecture significantly influenced species-level robustness, connectance, clustering coefficient and return time under two self-regulation parameterizations, whereas these effects generally weakened or disappeared after taxa were aggregated to the genus level. Reductions in taxonomic resolution can alter the estimated values of commonly used network metrics and consequently modify conclusions concerning network organization (Renaud et al., 2020; Rodrigues & Boscolo, 2020). Antagonistic interaction networks may be particularly sensitive to taxonomic aggregation, with loss of taxonomic resolution altering predicted network structure and some network descriptors (Llopis-Belenguer et al., 2023; Rodrigues & Boscolo, 2020). In the present system, aggregation at the genus level inevitably combines species that may differ in abundance, host use and trophic interactions. Aggregating such ecologically distinct species into a common node can conceal species-specific interaction patterns that are retained at finer taxonomic resolution (Llopis-Belenguer et al., 2023; Renaud et al., 2020). The loss of significant gall-architecture effects following aggregation therefore suggests that an important component of the structural differences between single- and multichambered communities occurs through species-specific interaction patterns within genera. Species-level identification may consequently be particularly important when ecological network analyses are used to investigate parasitoid communities associated with gall-inducing insects.

Gall architecture was also associated with differences in the relationship between community size and diversity. Although multichambered communities contained more taxa on average, Shannon diversity was significantly lower in multichambered networks after controlling for network size at both taxonomic resolutions. This indicates that the larger number of taxa occurring in multichambered communities does not necessarily translate into a more even distribution of abundance among community members. Instead, multichambered galls may accommodate a larger assemblage while remaining numerically dominated by a smaller subset of taxa (Bailey et al., 2009; Baine et al., 2024; Joseph et al., 2011; Rossi et al., 2006). The Gini index showed a pattern compatible with greater heterogeneity of interaction strength in multichambered communities, but the gall-architecture effect was not significant at either taxonomic resolution. Consequently, the lower Shannon diversity should not be interpreted as direct evidence for greater inequality in network interaction strength. Rather, the combined results suggest that gall architecture influences several different dimensions of community organization independently, with multichambered galls supporting larger and, at the species level, more highly connected and clustered networks, while their abundance distributions are comparatively less even. Such patterns are consistent with the broader view of gall morphology as an ecological filter that modifies the opportunities available to different members of the associated natural-enemy community rather than simply increasing parasitoid diversity (Bailey et al., 2009; Baine et al., 2024; Joseph et al., 2011; Stone & Schönrogge, 2003).

Taken together, our findings indicate that gall architecture is associated with measurable changes in the structure and stability of *Diplolepis*-associated parasitoid networks. The strongest evidence concerns structural stability, with multichambered networks showing greater robustness together with increased species-level connectance and clustering. Dynamical stability also differed between gall architectures, although this effect depended on the assumed strength of self-regulation and was detectable primarily at species-level resolution. In contrast, the predicted increase in Gini index was not statistically supported, while Shannon diversity indicated that larger multichambered communities were not necessarily more even. These results emphasize that ecological stability is multidimensional and that different components of stability need not respond identically to the same ecological driver (Donohue et al., 2016). More broadly, they suggest that the physical organization of the habitat generated by gall-inducing insects can influence ecological interactions beyond the direct gall inducer-parasitoid relationship and contribute to the emergent topology of the entire associated community (Bailey et al., 2009; Baine et al., 2024; Fang et al., 2024; Luz et al., 2021; Stone & Schönrogge, 2003). Our results further suggest that, in *Diplolepis*, these structural effects can extend to the stability of the resulting interaction network.

## Notes

### Competing Interest Statement

The authors have declared no competing interest.

